# Structural characterization of the *Pseudomonas Aeruginosa* MexR–*mexR* repressor-operator complex: a small-angle X-ray and neutron scattering perspective

**DOI:** 10.1101/2024.04.02.587325

**Authors:** Zuzanna Pietras, Francesca Caporaletti, Cy M. Jeffries, Vivian Morad, Björn Wallner, Anne Martel, Maria Sunnerhagen

## Abstract

The rapid spread of acquired multidrug resistance (MDR) in bacteria is a world-wide health threat. The MexR protein regulates the expression of the MexAB-OprM efflux pump, which actively extrudes chemical compounds with high toxicity to the host organism *Pseudomonas Aeruginosa*. In repression mode, two MexR dimers bind to an operator with two homologous pseudo-palindromic boxes located in proximity (named PI and PII). Here we report a first structural characterization of the complex in solution using small angle X-ray scattering (SAXS), small-angle neutron scattering (SANS) and rigid body modelling. The spacing between the PI and PII boxes is rich in AT base pairs indicate possible flexibility between the two MexR dimer binding sites. In agreement, our best modelling fits show a requirement for DNA bending between the two MexR binding sites to optimally fit SAS data as well as known biological properties of the MexR operons. Taken together, this study contributes to better understanding of the structural properties of bacterial operators and their repressor proteins.

## Introduction

Bacterium *Pseudomonas aeruginosa* belongs to a group of opportunistic pathogens associated with the debilitating nosocomial infections in immunocompromised individuals (1). It is infamous for being the main source of ventilator-associated pneumonia and chronic lung infection in cystic fibrosis patients (2). The effective treatments are particularly difficult to find due its high-level innate resistance to a vast range of antimicrobial agents and a quick adaptability to harsh environmental conditions. The antibiotic pressure is caused by a usage of combination therapies targeting multiple virulence mechanisms, resulting in the selection of diverse antibiotic-resistant strains (3). The factors of the inherent antibiotic resistance are contributed to a low outer membrane permeability and expression of proteins forming efflux systems which actively eliminate antibiotics out of the cell (4, 5). Several types of multidrug efflux pumps are encoded in *P. aeruginosa* genome, where MexAB-OprM and MexYX-OprM systems are already found in the wild-type strain and are highly overexpressed in hospital-identified variants (6–8).

The MexAB-OprM efflux pump shows a significant contribution to the antibiotic resistance, as it excretes the broadest spectrum of antibiotics, including chloramphenicol, β-lactams, quinolones, fluoroquinolones, novobiocin, and tetracycline (7). There are three transcription regulators that are showed to modulate expression of MexAB-OprM system (Fig. 1A). MexR is identified as a primary regulator that represses expression of *mexAB-oprM* and *mexR* genes by simultaneous binding to two operator sequences within *mexR-mexA* intergenomic region (9, 10).

**Figure 1.**
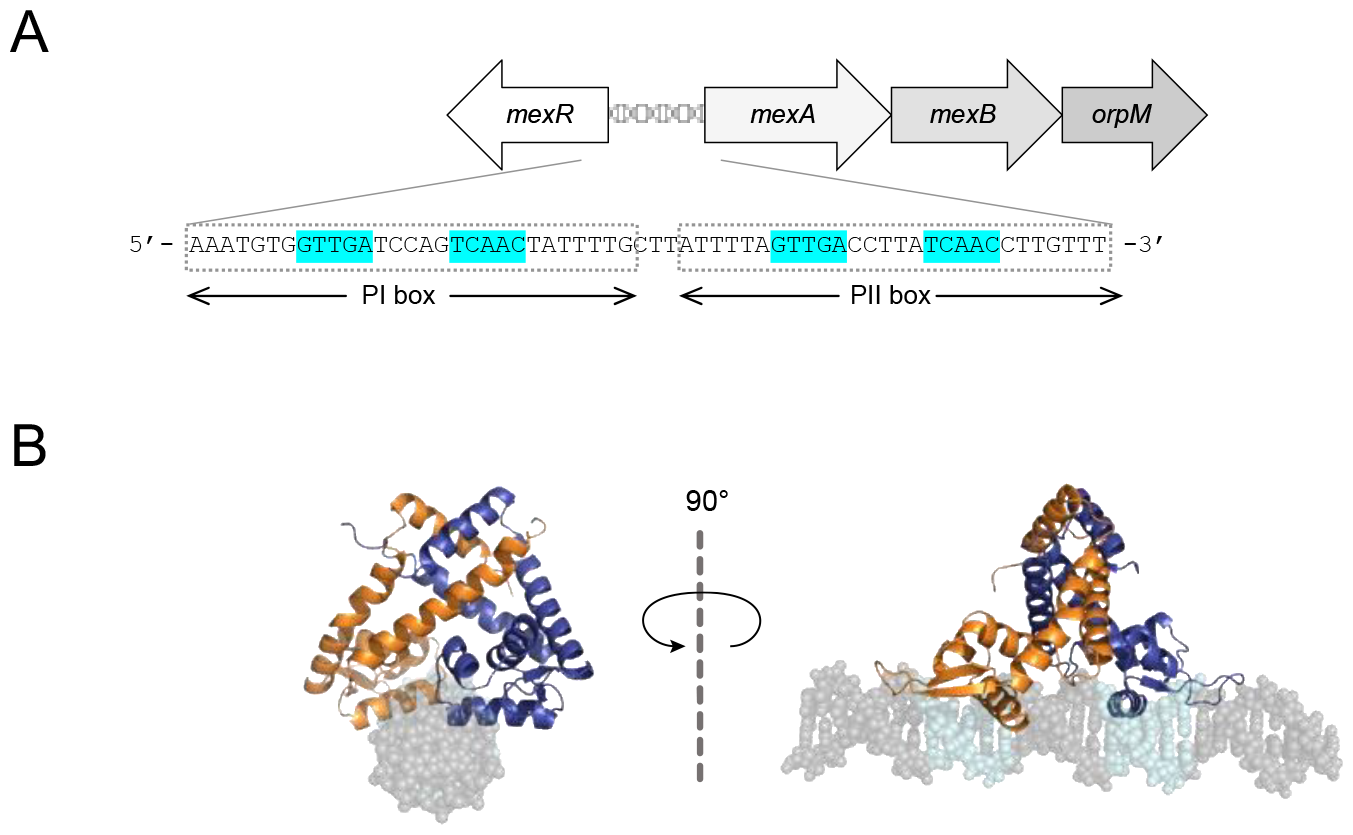
Schematic overview of the MexR transcription regulator and the promoter DNA sequences. **A**. The overview of MeAB-OprM operon and MexR promoter region. The DNA sequence of *mexR-mexA* intergenomic region with the MexR DNA recognition motifs highlighted in cyan. **B**. Model of a MexR dimer in complex with PII binding box.

MexR belongs to the MarR (multiple antibiotic resistance regulator) family of transcription regulators that actively control cellular response to harmful chemicals (11). It forms a pyramidal homodimer in solution adopting a pseudo-2-fold symmetry. Characteristic for this family is a winged helix-turn-helix (wHTH) motif that is essential for the DNA binding (9, 12). In the repression mode, the MexR recognition helices, α4-α4’, of each monomer are inserted into the consecutive major grooves whereas the antiparallel β-strand wings contact the closest minor grooves (Fig. 1B) (13). In the previous studies, Dnase I footprinting experiments identified the exact DNA recognition sequences of MexR binding regions (9, 10). It has been shown that MexR recognises two regions, named PI and PII, of approximately 28 base pairs separated by 3 unprotected bp (9). Each contains a pair of inverted palindromic GTTGA sequences partitioned by 5 bp, which would predict MexR binding on the same face of the DNA, providing linear B-DNA in solution. However, no structures of MexR-DNA complexes have been solved to date, limiting our understanding of the structural mechanisms in repression.

Over the decades, a significant number of biochemical and structural data of multiple members of MarR family have been reported (11, 14, 15). Very few of these structures comprise MarR-family - DNA complexes (16–26) and only one shows MarR family repressors bound to adjacent DNA binding sites (27). Recent high-resolution structures of bacterial transcription regulators in complex with two binding DNA boxes indicate that the spacing between such boxes is both structurally and functionally important (28–30). Extrapolating from this, the variability within the organisation of the intergenic regions of bacterial operons suggests that each MarR homologue will direct its own signature change in the DNA topology upon binding (11). The distance between the DNA boxes repressors in such bacterial operons, as well as its sequence, may also contribute to such signature changes and thereby be part of the binding mechanism as well as its functional implications.

In our previous work, we have reported *ab initio* and molecular-based models of MexR bound to one of the DNA boxes (PII), based on small angle x-ray and neutron scattering data and molecular modelling. Together, this suggested a DNA-binding conformational selection mechanism, where MexR in the absence of DNA is highly flexible, presenting a wide ensemble of conformations, where some are equivalent to those observed for the bound state (Caporaletti *et al*.).

Here we report a low-resolution structure of MexR in complex with its two adjacent binding regions, derived from small angle x-ray and neutron scattering data. The *ab initio* model shows similar conformation features to a complementary rigid body model, both presenting an elongated dumbbell structure with two protein species placed asymmetrically to each other.

## Materials and methods

### Protein expression and purification

The gene encoding MexR (Val5-Leu139) from *P. aeruginosa* was cloned into pNH-TrxT expression vector (31) with N-terminal His_6_-Thioredoxin (Trx) tag and Tobacco Etch Virus (TEV) protease cleavage site. Heat-shock transformed *Escherichia coli* Rosetta 2(DE3) cells were grown in LB media containing 50 µg/mL kanamycin and 34 µg/mL chloramphenicol at 37 ºC to OD_600_ of 0.9. Culture was induced with 0.5 mM isopropyl β-d-1-thiogalactopyranoside (IPTG) and shaken at 18 ºC for overnight protein expression. Cells were harvested through centrifugation and lysed by sonication in Buffer A (50 mM sodium phosphate, 300 mM NaCl, 10 mM Imidazole, 5% v/v glycerol, 10 mM β-mercaptoethanol, pH 7.1) supplemented with 5 µg/mL DNase I (Sigma-Aldrich) and cOmplete protease inhibitors cocktail tablet (Roche).The lysate was cleared by centrifugation at 20 000 x g for 30 minutes and the supernatant was loaded onto a Ni-NTA resin equilibrated in Buffer A. After 1 h incubation at 4 ºC, column was extensively washed with Buffer A followed by a wash with Buffer B (Buffer A with 30 mM imidazole). His_6_-Trx-MexR was eluted in Buffer C (Buffer A with 300 mM imidazole) and TEV protease was added to the eluent prior to dialysis against Buffer A at 4 ºC overnight. The protein mixture was incubated for 30 minutes with Ni-NTA resin pre-equilibrated in Buffer A for the removal of uncleaved species and free His_6_-Trx tag. The collected flow-through was subjected to the final purification step by gel filtration using a HiLoad 16/600 Superdex 75 pg column (Cytiva) equilibrated in 20 mM HEPES, 150 mM NaCl, 5% v/v glycerol, 2 mM TCEP. Each purification step was analysed by SDS-PAGE (Bio-Rad) with Coomassie Blue stain to assess the protein purity. The protein concentration was determined using Bradford protein assay (32) with the absorbance measured at 595 nm using microplate reader (Tecan).

To express partially deuterated MexR, *E. coli* Rosetta 2(DE3) cells containing pNH-TrxT-MexR were grown on agar plates supplemented with appropriate antibiotics at 37 ºC overnight. The overnight colonies were used to inoculate 2 litres of M9 minimum media (6 g/L Na_2_HPO_4_, 3 g/L KH_2_PO_4_, 0.5 g/L NaCl, 2 g/L NH_4_Cl, 0.1 mM CaCl_2_, 1 mM MgSO_4_, 10 mg/mL biotin, 1 mg/mL thiamine) containing 99.9% D_2_O (Sigma-Aldrich) and unlabelled glycose (4 g/L). Cells were grown 37 ºC to OD_600_ of 0.8 and 0.5 mM IPTG dissolved in D_2_O was added to induce protein expression. The temperature was reduced to 20 ºC and cultures were incubated at 100 rpm overnight. Deuterated protein was purified in the same manner as described above using hydrogenated buffers. The non-exchangeable average deuteration level was determined by MALDI-TOF (Bruker) using comparison of purified unlabelled and labelled samples. The calculated deuteration level was 72 % and the deuterated MexR is called dMexR through the text.

### Preparation of double-stranded DNA and MexR-DNA complex

Complementary single-stranded oligonucleotides were purchased (Eurofins Genomics) and contain the PI and PII sites based on the physiological operator sequence (9): 5’-AAATGTG**GTTGA**TCCAG**TCAAC**TATTTTGCTTATTTTA**GTTGA**C CTTA**TCAAC**CTTGTTT-3’ (palindromes are in bold). The ssDNAs were resuspended in annealing buffer (20 mM Tris-HCl, pH 8.0, 150 mM NaCl, 1 mM EDTA). Equimolar concentrations of complementary strands were mixed and incubated at 95ºC for five minutes and slowly cooled down to room temperature. Purified protein was mixed with dsDNA keeping a 4:1 ratio and incubated for 1 hour at room temperature. The formed complex was further purified by size exclusion chromatography (SEC) with a Superdex 200 Increase 10/300 column (Cytiva) equilibrated in 20 mM HEPES, 150 mM NaCl, 2 mM TCEP.

### Size exclusion chromatography coupled to multi-angle light scattering (SEC-MALS)

The MALS data were measured using Mini-Dawn TREOS coupled to an OptiLab T-Rex refractometer (Wyatt Technologies). The experiments were performed at 20ºC and 30 μL of sample was injected into a Superdex 200 Increase 5/150 column (Cytiva) equilibrated in 20 mM HEPES, 150 mM NaCl, 1% glycerol, 2 mM TCEP, at a flow rate of 0.3 mL/min. The molecular weight (MW) distribution of the species eluting from the column was estimated from the MALS intensities and concentration estimates obtained from a refractive index (RI). The input values of the differential refractive index increment (dn/dc) were 0.186 mL/g and 0.175 mL/g (33) for protein and dsDNAs respectively and 0.180 mL/g for MexR-DNA complexes. The component analysis of the complexes was performed using Conjugate Analysis algorithm from ASTRA 6 Software package (Wyatt Technologies).

### Isothermal titration calorimetry

Isothermal titration calorimetry measurements were performed on a MicroCal PEAQ-ITC instrument (Malvern). Beforehand, protein and dsDNAs were extensively dialyzed against buffer containing 20 mM NaPO_4_, pH 7.0, 150 mM NaCl, 5 mM β-mercaptoethanol and centrifuged at 20 ºC at 3500 rpm for 10 minutes. Binding experiments were done with 105 μM protein solution in the syringe and 2 μM dsDNA in the cell. The titration started with an initial injection of 0.4 μL and was followed by 18 consecutive 2.3 μL injections into sample cell at 20 ºC. Analysis of the data were performed using MicroCal PEAQ-ITC Analysis Software (Malvern) according to the one (two) binding site model.

### Small angle scattering data collection

Small angle X-ray scattering data were measured at the EMBL BioSAXS-P12 beamline (34) utilising an in-line SEC-SAXS setup. The used column and elution parameters were the same as described above for the SEC-MALS and experimental details, including injection volumes and concentrations of each sample are listed in Table S1. Automated sample injection and data collection were controlled using the BECQUEREL beam line control software (35). The SAXS intensities were measured from the continuously flowing column eluent as 2400 x 1 s individual X-ray exposures using a Pilatus 6M detector. The 2D-to-1D data reduction, i.e., radial averaging of the data to produce 1D, I(*q*) vs *q* profiles, were performed using the SASFLOW pipeline (35, 36) incorporating RADAVER from the ATSAS 3.0 suite of software tools (37). The 2400 individual frames obtained for each SEC-SAXS run were processed using CHROMIXS (38). Briefly, individual SAXS data frames were selected across the SEC-elution peak of the protein. An appropriate region of the elution profile, corresponding to SAXS data measured from the solute-free buffer, were identified, averaged, and then subtracted from the SEC-peak frames to generate individual background-subtracted sample data frames. These individual data frames underwent further CHROMIXS analysis, including the assessment of the radius of gyration (R_g_) of the sample through the SEC peak. Then the individual frames were scaled (to consider changes in concentration) and averaged to produce a final 1D-reduced and background-corrected scattering profile. Only those SAXS data frames with a consistent R_g_ through the SEC-elution peak and evaluated as statistically similar through the measured q-range (0.024–7.3 nm^-1^) were used to generate the final SAXS profiles.

Small angle neutron scattering data were collected on the D22 instrument at the Institute Laue Langevin (ILL). Two aliquots of each MexR-DNA complex at ∼ 6 mg/mL were dialysed in 20 mM HEPES, 150 mM NaCl and 2 mM TCEP buffer, one against buffer containing 0% D_2_O contents and the second with 100% D_2_O. The two-dimensional data were reduced to 1-D SANS-profiles using GRASP software (39) and scattering profiles from the two detector positions were merged and background-subtracted using PRIMUS (40).

### Small angle scattering data analysis and modelling

Data analyses were conducted using the ATSAS package and the determination of the radii of gyration, R_g,_ and forward scattering intensities, *I*(0), were extracted using Guinier approximation (ln*I*(q) versus q^2^, for qR_g_ < 1.1) (41). The GNOM software (42) was used to calculate the pair-distance distribution function, p(r), and to estimate maximum particle dimensions, D_max_. The ab initio shape reconstruction of dsDNAs was performed using program DAMMIF (43). Ten individual models were generated, and their consistency was evaluated using the normalized spatial discrepancy (NSD) metric and processed using DAMAVER set of programs (44) to produce a final dummy atom model representing consistent structural features. Rigid-body modelling using program SASREF (45) was used to refine solution conformations of MexR-DNA complex against SAXS data. The models were constructed by combining a high-resolution structure of MexR (PDB: 1LNW) (12) and a B-DNA model derived using 3D-DART server (46). The B-DNA model was divided into five segments: 5’-AAATGTG**GTTGA**TCCAG**TCAAC**T ^*^ ATTTT ^*^ GCTT ^*^ ATTTT ^*^ A**GTTGA**CCTTA**TCAAC**CTTGTTT-3’; palindromes are in bold and star ^*^ indicates a breakpoint; with a distance restrain of 7 Å. The final fits to the experimental SAXS and SANS data were computed using CRYSOL (37) and CRYSON (47) and evaluated using the reduced χ^2^ test (48). The details about SAS data collection and processing are presented in Supplemental Table 1 and Supplemental Table 2.

## Results

### Stoichiometry and affinity of the MexR-DNA complex

To characterize the interactions of MexR with DNA, we performed Isothermal Titration Calorimetry experiments (ITC). Titration of MexR to the 60 bp double-stranded DNA comprising PI and PII boxes resulted in formation of a stable heterocomplex with a dissociation constant of 206 ± 52 nM (Figure 2A). The MexR binds to the two promoter regions with similar affinities as for individual boxes. The previously reported Kd value from Surface Plasmon Resonance measurements for MexR-PI interactions is 370 nM (49), while ITC experiments of MexR-PII binding indicated dissociation constant of 240 nM (50). The stoichiometry of the complex implies binding of four MexR monomers to one dsDNA, which is further confirmed by SEC-MALS-RI (Fig. 2B). The complex eluted from the column as a single peak and the calculated Molecular Weight (MW) is within the range of 97.4-99.8 kDa, with an average of 98 kDa. The expected MW of the complex calculated from the amino acid and nucleic acid sequence is 101kDa, assuming 4:1 protein:dsDNA ratio. The Protein conjugate analysis indicates the average MW of protein fractions is 58.5 kDa (63.3 kDa is a calculated MW of four MexR monomers), while the modifier, dsDNA, is around 41.6 kDa (36.8 kDa based on the nucleotide sequence) (Fig. 2B). Taken together, the data indicates that the whole complex has the average MW of 100 kDa, what corresponds well with the theoretical MW of 100 kDa.

**Figure 2.**
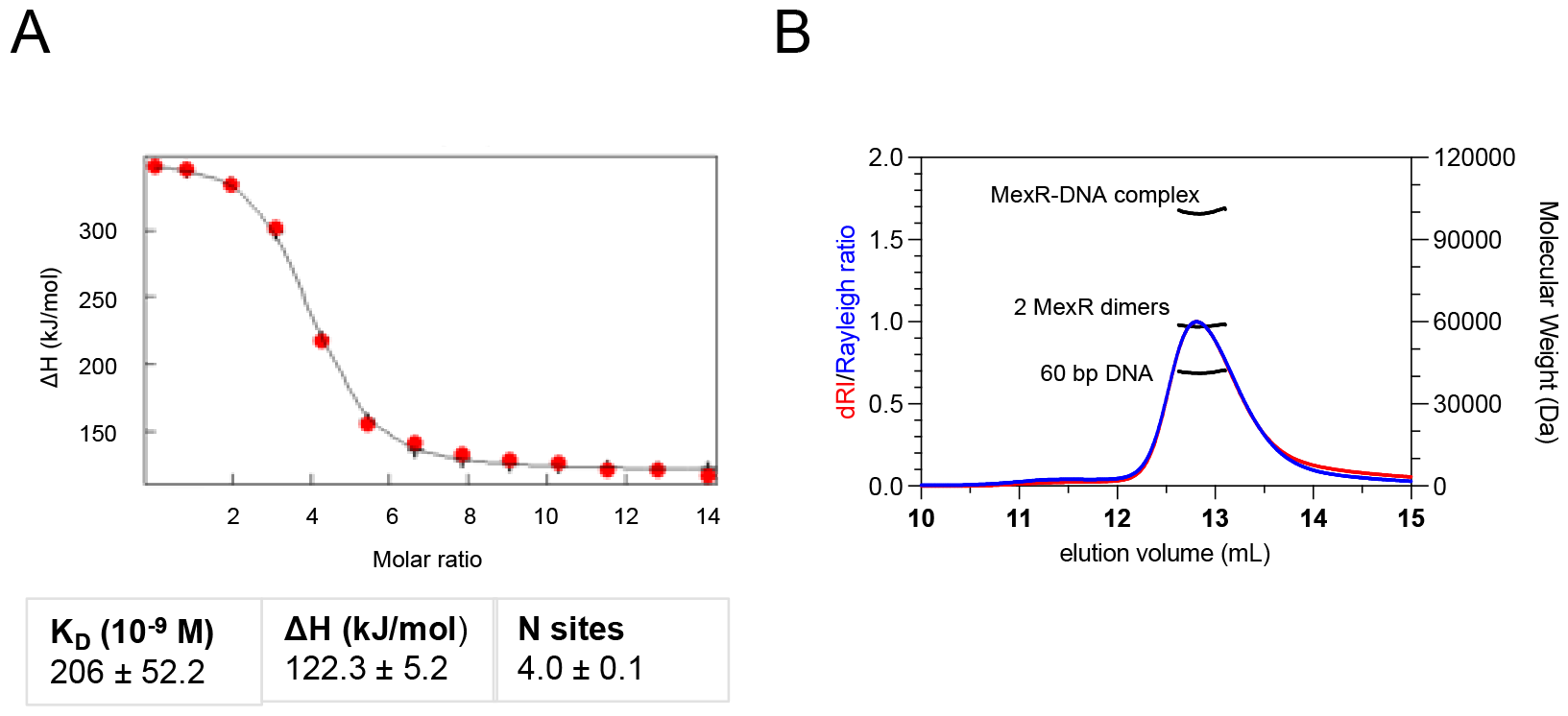
Two MexR homodimers bind to *mexR-mexA* intergenomic region. **A**. Binding curve and fit of ITC measurements of MexR titrated to 60 bp double-stranded DNA comprising two pairs of inverted palindromic boxes, PI and PII. Data was fitted using one set of sites model. **B**. SEC-MALS-RI elution profile and calculated molecular mass of the complex and conjugation analysis to determine quantities of individual components.

### Small angle X-ray scattering data suggests an extended, slightly bent complex

SEC-SAXS experiments were performed to investigate the overall shapes of the dsDNA alone and the effects of MexR-DNA complex formation. The final SAXS profiles obtained from the scaled and averaged individual data frames is indicated in Figure 3A. The radii of gyration, *R*_*g*_, were derived using Guinier analysis (within the limit *qR*_*g*_ < 1.1), as 5.14 and 5.20 nm for DNA and the complex respectively (Table 1 and Supplemental Table 1). The DNA shows the characteristics of an extended rod-like particle (Fig. 3A) allowing to extract *R*_*g*_ of cross-section with stable value of 0.84 nm. Dimensionless Kratky plots of SAXS profiles, further confirms that both DNA alone and the MexR-DNA complex are extended in solution (Fig. 3B). The appearance of the *p*(*r*) profile suggests a rod-like particle of the dsDNA in solution, while the complex shows features of a dumbbell with a maximum at 4 nm and a secondary shoulder at 10 nm (Fig. 3C). The *R*_*g*_ of dsDNA, determined from *p*(*r*) analysis of 5.36 nm is consistent with the Guinier *R*_*g*_ estimates, while the *D*_*max*_ is evaluated at 19 nm (Table 1, Fig. 3C). The *D*_*max*_ of the complex is estimated to be around 18.5 nm, suggesting that both particles are similar in size (Fig. 3C) and the *R*_*g*_ derived from the *p(r)* of 5.38 nm complementing the Guinier estimates (Table 1). A fully extended DNA of this size has an end-to-end distance of 20.2 nm. Combined, the structural parameters suggest that MexR protein likely decorates a predominantly extended, possibly very slightly bent, DNA molecule, that may undergo further slight conformational adjustments on engaging with the protein.

**Table 1.**
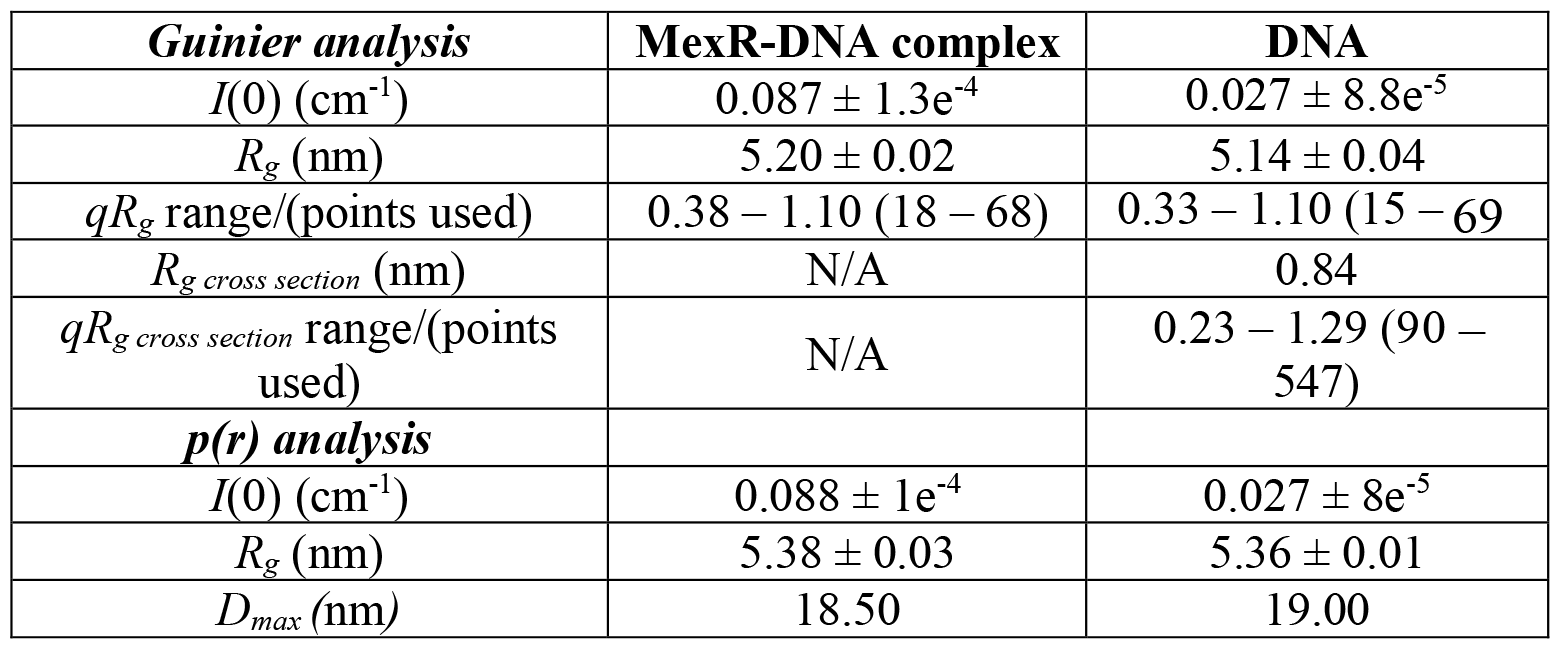
Structural parameters derived from the experimental SAXS data.

**Figure 3.**
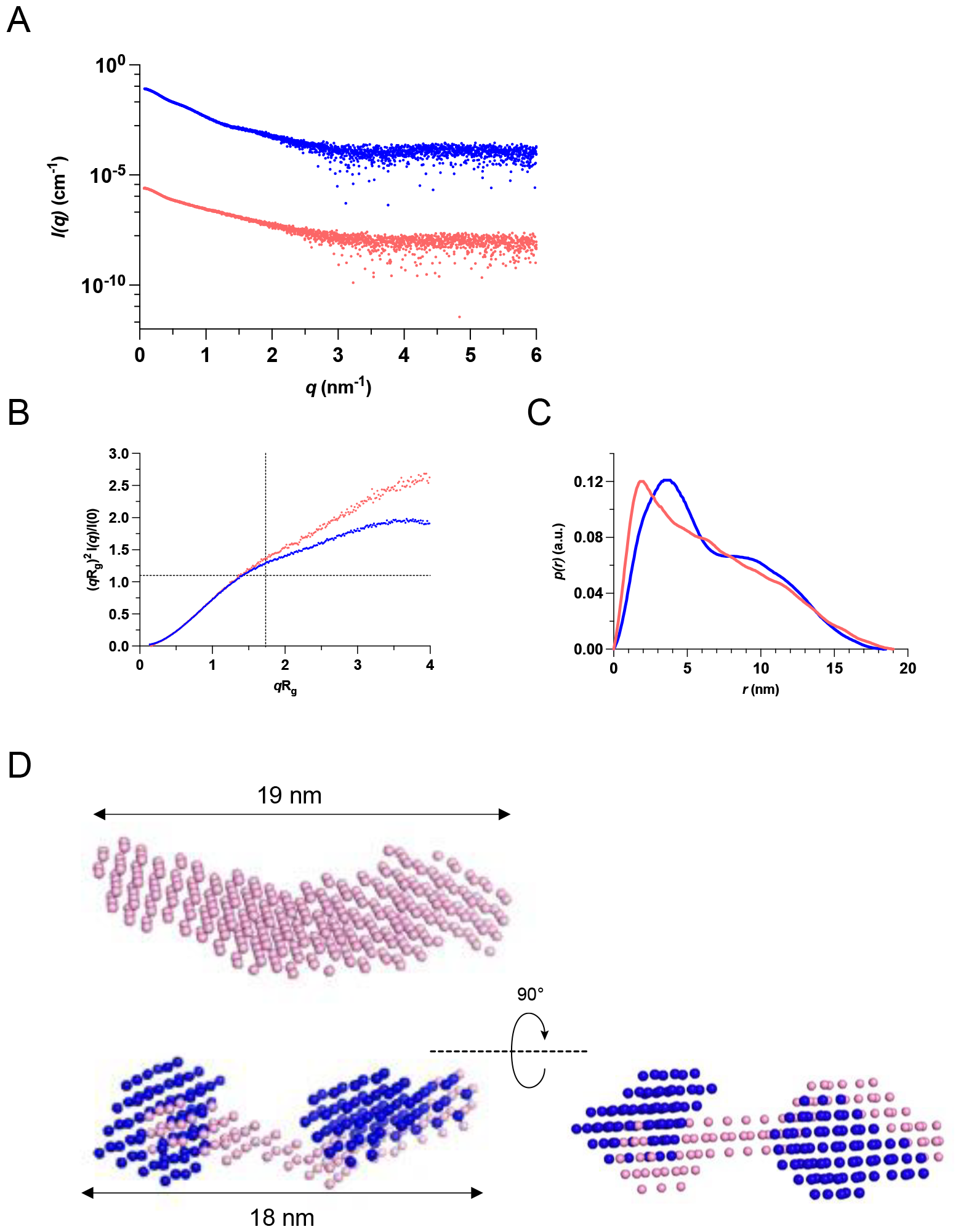
The solution conformations of MexR-DNA complex and DNA alone. **A**. The SAXS scattering curves collected from MexR-DNA complex (blue) and 60 bp DNA (pink). Offset for clarity. **B**. Dimensionless Kratky plot derived from SAXS data indicates that both macromolecules are extended in solution. **C**. The pair distribution functions p(r) indicates D_max_ for the complex (18.5 nm) and DNA (19 nm). **D**. Low resolution envelope of DNA derived using DAMMIF and MONSA model of MexR-DNA complex, where MexR dimers are in blue, and DNA is coloured pink.

In agreement with the model-free analysis, a derived ab initio DAMMIF model of dsDNA shows an extended envelope with a slight bend (Fig. 3D). A low-resolution multi-phase MONSA model of MexR-DNA suggest that two MexR homodimers are positioned on the DNA at distinct, non-interacting positions (Fig. 3D).

### Molecular modelling suggests DNA flexibility between binding sites in the MexR-DNA complex

To explore structural features of the DNA binding mechanism of two MexR homodimers, three different approaches were performed and compared based on a goodness-of-fit test for SAXS data (Fig. 4A). The first model, Model 1, was built by manual docking of MexR on the PI and PII boxes, based on a homology OhrR-*ohrA* operator complex (16) as a template, and with the B-DNA structure generated as a straight model. The wHTH domains are then positioned into the DNA major grooves within the inverted palindromic GTTGA sequence, resulting in a model where the MexR proteins are placed on the same face of the DNA (Fig. 4B). The model was evaluated using CRYSOL and fitted to the experiment SAXS data, showing a poor fit at low angles (Fig. 4A) and a χ^2^ of 6.01. To further improve the fit, SASREF software was used, where the MexR homodimers were allowed to refine their position on the DNA against the SAXS data without any positional constraints. Compared to Model 1, in this Model 2, the MexR dimers move towards each other, away from their designated binding motifs, which one of the dimers has entirely released, and the proteins are no longer positioned on the same face of the DNA (Fig. 4B). Interestingly, while this placement of the dimers significantly improved fit to the SAXS data, reducing the χ^2^ to 1.01, this model is inconsistent with previous knowledge of the MexR binding sites (9).

**Figure 4.**
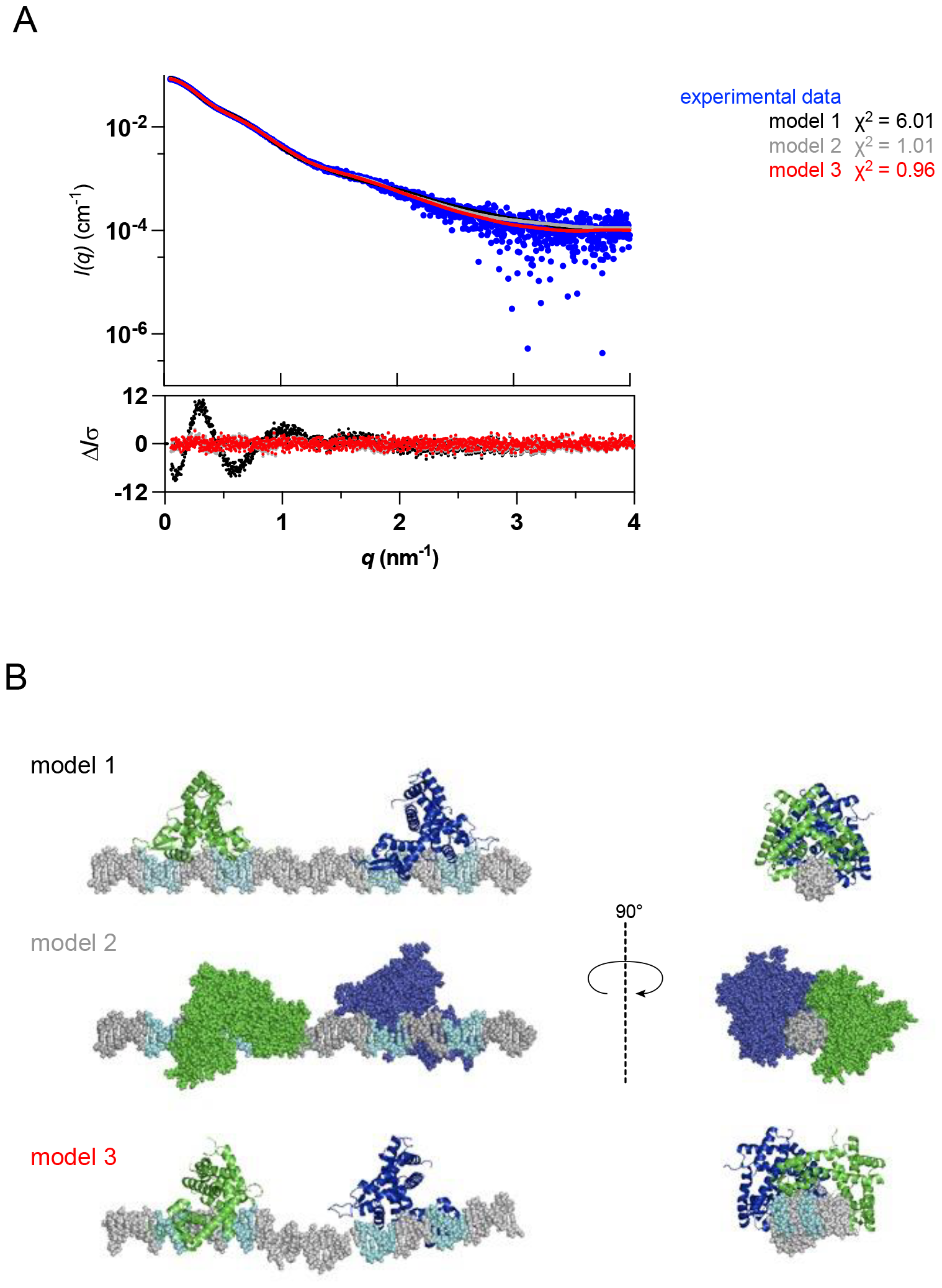
Comparison of three molecular models of two MexR dimers bound to the promoter DNA. **A**. The SAXS profile and calculated fits of the models. **B**. Models of the MexR-DNA complexes, the structures are rotated 90º along y axis to visualize different positioning of MexR dimers on the DNA.

To allow change of the relative position between the bound MexR dimers with maintained anchoring to the inverted repeats, a third model was derived allowing for flexibility in the AT-rich linker between the PI and PI binding sites. Here, the DNA model was portioned into five pieces, maintaining the MexR dimer binding to the promoter regions as described in (50), and with contact points designed to place the DNA segments according to the correct bp sequence within a distance of 5 Å. After SASREF refinement against SAXS data, the resulting third model (Model 3) displays a similar positioning of the proteins as in the second model (Fig. 4B). The main difference is visible in the overall shape of the DNA, where we can observe an approximately 30° bend in the linker region between the PI and PII DNA sites, and a slight tilting of DNA towards one side. Model 3 also displays a good fit to SAXS data, with the χ^2^ of 0.96. To further distinguish whether one of the models can be a more accurate representation of the complex, we conducted a SANS experiment. The two theoretical models (2 and 3) were fitted to the neutron contrast variation data set (Fig. 5). The obtained χ^2^ values suggest that both molecular models reasonably fit the data and display similar trends and discrepancies at each percentage of D_2_O (Supplemental Table 2). However, while the assessment of the data using AMBIMETER (51) shows that while the 3D reconstruction of the DNA alone can be potentially unique (ambiguity score of 1.08), the shape topology of the complex might be ambiguous (ambiguity score of 2.5). Taken together, this suggests that a full description of the MexR-DNA complex in solution studied here may require an ensemble of conformations with different propensities to properly reflect the combined SAS data.

**Figure 5.**
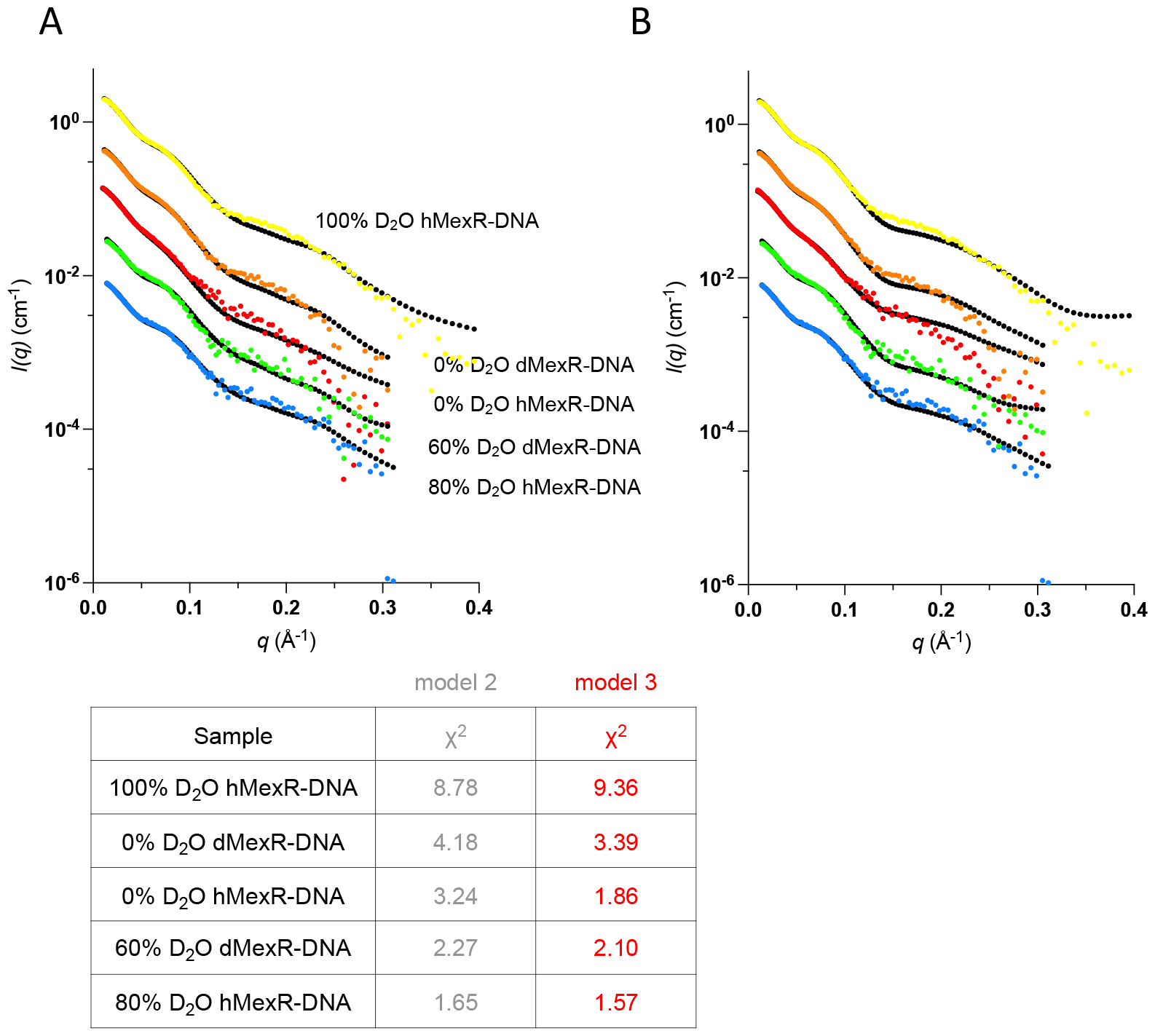
SANS data collected on hydrogenated and deuterated MexR in complex with the DNA. **A**. CRYSON fits (black dotted line) to the experimental SANS data for model 2. **B**. CRYSON fits (black dotted line) to the experimental SANS data for model 3. The χ2 values for each sample are display in the table.

## Discussion

In this work, we have employed a small angle scattering approach to study the envelope of MexR dimers bound to its operator site in the Pseudomonas Aeruginosa genome. In a recent study, we show by SAS that in solution MexR binds to a mexR palindrome as would be expected, but conformationally selecting an asymmetric binding mode in the bound state that shifts the position of the two MexR dimers from being at the same side to being at a slight angle towards each other (50) MexR belongs to the large MarR family of homodimeric bacterial repressors (15, 52), where crystal structures of DNA complexes are still very scarce but still steadily increasing in numbers (16–26). In all of these complex structures, the bound DNA is straight, showing only small variations from B-DNA. Only one of these crystal structures represents two dimer repressors bound to the same operator (27), even if multiple binding sites on DNA are commonly observed for this family of proteins (11). This protein-DNA complex comprises two MarR family proteins bound on opposite sides of a fairly straight 30-mer B-DNA, and where the two MarR proteins are close enough to share the same major groove (27).

The present study is the first direct observation of the structural envelope of a MarR family protein bound to an extended operator sequence containing two MarR protein binding sites. In the extended *mexR* operator, palindrome positioning suggests that MexR protein dimers would bind on the same face of an assumed straight, B-DNA sequence. However, we show that this model fits scattering data poorly; rather, the MexR dimers appear to be closer to each other in the bound state. Good fits to data were obtained either in B-DNA models where MexR dimers were positioned closer to each other than the palindromes would suggest (Model 2), or where the DNA linker between the palindrome needs to be structurally remodeled to a slightly bent conformation (Model 3). While the agreement in relative positioning of the MarR dimers relative to the DNA axis in Models 2 and 3 is impressively similar, only the bent model (Model 3) agrees with appropriate positioning of MexR dimers onto the corresponding palindromes (9) as expected from previously determined MarR family DNA complexes (16–26). Our analysis suggests that a single structure is not sufficient to describe how two MexR binds to its full operator. It is possible or even likely that a full ensemble describing the MexR-DNA complex studied here would comprise a fan of variously bent states, with the best single model shown here only being a representative member of such an ensemble. Further data analysis and experimental input from, for example, molecular dynamics simulations and NMR (53) will be required to analyze these dynamic aspects further.

Specific DNA recognition by proteins is fundamental to the regulation of processes controlling all aspects of life and is intrinsically a highly dynamic process (54). Our view of protein-DNA complexes has been strongly influenced crystallization, where DNA often stabilizes the gitter by end-to-end contacts, resulting in extended DNA helix patterns throughout the crystals (55). This favorably biases structure determination of protein-DNA complexes with fairly straight DNAs, which is also clearly reflected in the PDB database. In contrast, protein-DNA complexes that hold inherent flexibility, show asymmetric properties and/or form dynamic ensembles therefore present challenging targets of study.

We show here that small-angle scattering can evaluate and resolve the envelope of such complexes, without imposing the restrictive environment of the crystal (53). The inability to accurately represent this critical group of dynamic complexes by crystallography alone provides an ample window for exploration by small-angle scattering in solution (56) together with NMR and cryo-EM, with the aim to depict such ensembles in full dynamic and structural detail (57).

