## Supplemental Table 1 for "Structural characterization of the *Pseudomonas Aeruginosa* MexR–*mexR* repressor-operator complex: a small-angle X-ray and neutron scattering perspective"

| **Instrument Parameters** | |
| --- | --- |
| Instrument | EMBL-P12 bioSAXS beamline with Dectris Pilatus 6M detector |
| Wavelength (nm) | 0.124 |
| Sample to detector distance (m) | 3 |
| Measurement type | SEC-SAXS |
| Column | Superdex 200 Increase 5/150 |
| Injection volume (µL) | 30 |
| Flow rate (mL/min) | 0.3 |
| Solvent | 20mM HEPES, pH 7.1, 150mM NaCl, 10mM DTT, 1% v/v glycerol |
| Exposure time (s)/number of frames | 1/2880 |
| *q* measured range (nm^-1^) | 0.0024 – 0.73 |
| Temperature (^o^C) | 20 |
| **Software employed for SAXS data reduction and analysis** | |
| SAXS data reduction | SASFLOW |
| Extinction coefficient estimate | ProtParam |
| Calculation of Δρ and v | MULCh |
| Structural analysis | PRIMUSqt, GNOM (ATSAS) |
| Shape/bead modelling | DAMMIF, MONSA |
| Atomistic modelling | SASREF/SASREFCV |
| Three-dimensional graphic representation | PyMOL |
| **Sequences** | |
| MexR (UniProt ID: P52003) residues 5 – 139 | SM – VNPDLMPALMAVFQHVRTRIQSELDCQRLDLTPPDVHVLKLIDE  QRGLNLQDLGRQMCRDKALITRKIRELEGRNLVRRERNPSDQRSFQL  FLTDEGLAIHQHAEAIMSRVHDELFAPLTPVEQATLVHLLDQCL |
| 60 bp DNA | 5’ – AAATGTGGTTGATCCAGTCAACTATTTT  GCTTATTTTAGTTGACCTTATCAACCTTGTTT – 3’ |

| **Sample details** | DNA short |
| --- | --- |
| Concentration (mg/mL) | 3.00 |
| Partial specific volume, *v* (cm^3^ g^-1^) | 0.592 |
| Particle contrast, Δρ (10^10^cm^-2^) | 5.319 |
| **Information content** | |
| #Shannon channels | 40 |
| Highest useable q_max_ (nm ^-1^) | 6.72 |
| Predicted *D_max_* (nm) | 18.7 |
| **Guinier analysis** | |
| *I*(0) (cm^-1^) | 0.0270 ± 0.0001 |
| *R_g_*, (nm) | 5.14 ± 0.04 |
| *qR_g_* range / (points used) | 0.33 – 1.10  (15 – 69) |
| *R_g cross section_* (Rod-Guinier, nm) | 0.84 |
| *qR_g cross section_* range / (points used) | 0.23 – 1.29  (90 – 547) |
| ***p(r*) analysis** | |
| *I*(0) (cm^-1^) | 0.0268 ± 0.0001 |
| *R_g_* (nm) | 5.36 ± 0.03 |
| *D_max_ (*nm*)* | 19 |
| Total estimate from GNOM | 0.62 |
| Quality of fit (χ^2^ / P-value) | 0.96 / 0.52 |
| Porod volume (nm^3^) | 88 |
| **Molecular Weight analysis** | |
| Theoretical MW | 45.6 |
| MW from SEC – MALLS (kDa) | 53 |
| MW Vc (kDa) | 41 |
| MW DARA (kDa) | 46 |
| **Shape classification** | |
| Classification / (predicted *D_max_*, nm) | Extended (17.9) |
| Ambimeter score (q*R_g_^max^*) | 1.1 |
| Number of shape topologies | 12 |
| Uniqueness | potentially unique |

| **Sample details** | MexR-DNA |
| --- | --- |
| Concentration (mg/mL) | 6.70 |
| Partial specific volume, *v* (cm^3^ g^-1^) | 0.686 |
| Particle contrast, Δρ (10^10^cm^-2^) | 3.575 |
| **Information content** | |
| #Shannon channels | 40 |
| Highest useable q_max_ (nm ^-1^) | 6.73 |
| Predicted *D_max_* (nm) | 18.7 |
| **Guinier analysis** | |
| *I*(0) (cm^-1^) | 0.0870 ± 0.0001 |
| *R_g_*, (nm) | 5.22 ± 0.02 |
| *qR_g_* range / (points used) | 0.38 – 1.10  (18 – 68) |
| *R_g cross section_* (Rod-Guinier, nm) | N/A |
| *qR_g cross section_* range / (points used) | N/A |
| ***p*(*r*) analysis** | |
| *I*(0) (cm^-1^) | 0.0880 ± 0.0001 |
| *R_g_* (nm) | 5.38 ± 0.01 |
| *D_max_ (*nm*)* | 18.5 |
| Total estimate from GNOM | 0.73 |
| Quality of fit (χ^2^ / P-value) | 0.96 / 0.30 |
| Porod volume (nm^3^) | 157 |
| **Molecular Weight analysis** | |
| Theoretical MW (kDa) | 106 |
| MW from SEC – MALLS (kDa) | 98 |
| MW Credibility Interval (kDa) | 114 |
| MW SAXSMow (kDa) | 105 |
| MW Vc (kDa) | 87 |
| **Shape classification** | |
| Classification / (predicted *D_max_*, nm) | Random-chain (N/A) |
| Ambimeter score (q*R_g_^max^*) | 2.50 |
| Number of shape topologies | 313 |
| Uniqueness | Might be ambiguous |
