## Supplemental Table 2 for "Structural characterization of the *Pseudomonas Aeruginosa* MexR–*mexR* repressor-operator complex: a small-angle X-ray and neutron scattering perspective"

| **Instrument Parameters** | |
| --- | --- |
| Instrument | D22 SANS beamline at ILL (Grenoble, France) |
| Wavelength (A) | 6 ± 0.6 |
| Detector distances (m) | 5.6 : 5.6 / 1.6 : 2.8 |
| Measurement type | Cuvette SANS |
| Absolute scaling method | Incident beam flux |
| Normalization | Divided by sample concentration |
| *q* measured range (A^-1^) | 0.014 – 0.5 |
| Solvent | 20 mM HEPES, pH 7.1, 150 mM NaCl, 2 mM TCEP |
| Temperature (^o^C) | 10 |
| **Software employed for SANS data reduction and analysis** | |
| SANS data reduction | GRASP |
| Extinction coefficient estimate | ProtParam |
| Calculation of Δρ and v | MULCh |
| Structural analysis | PRIMUSqt, GNOM (ATSAS) |
| Shape/bead modelling | MONSA |
| Atomistic modelling | SASREF/SASREFCV |
| Three-dimensional graphic representation | PyMOL |
| **Sequences** | |
| MexR (UniProt ID: P52003) residues 5 – 139 | SM – VNPDLMPALMAVFQHVRTRIQSELDCQRLDLTPPDVHVLKLIDE  QRGLNLQDLGRQMCRDKALITRKIRELEGRNLVRRERNPSDQRSFQL  FLTDEGLAIHQHAEAIMSRVHDELFAPLTPVEQATLVHLLDQCL |
| 60 bp DNA | 5’ – AAATGTGGTTGATCCAGTCAACTATTTT  GCTTATTTTAGTTGACCTTATCAACCTTGTTT – 3’ |
| 74 bp DNA | 5’ – GGGGTAAATGTGGTTGATCCAGTCAACTATTTT  GCTTATTTTAGTTGACCTTATCAACCTTGTTTCAGGTCCCC – 3’ |

| **Sample details** | **MexR – 60bp DNA complex** | | | | |
| --- | --- | --- | --- | --- | --- |
|  | dMexR-DNA  0% D_2_O | dMexR-DNA  55% D_2_O | dMexR-DNA  90% D_2_O | hMexR-DNA  0% D_2_O | hMexR-DNA  78% D_2_O |
| Concentration (mg/mL) | 6.37 | 6.44 | 6.45 | 6.82 | 7.04 |
| Particle contrast, Δρ (10^10^cm^-2^) | 5.071 | 1.881 | - 0.155 | 2.792 | - 1.732 |
| MexR contrast, Δρ (10^10^cm^-2^) | 5.678 | 2.610 | 0.658 | 2.341 | - 2.010 |
| DNA contrast, Δρ (10^10^cm^-2^) | 3.763 | 0.297 | - 1.908 | 3.763 | - 1.152 |
| Partial specific volume, *v* (cm^3^ g^-1^) | 0.685 | | | | |
| **Guinier analysis** | | | | | |
| *I*(0) (cm^-1^) | 0.4600 ± 0.002 | 0.0620 ± 0.001 | 0.0063 ± 0.0001 | 0.2900 ± 0.002 | 0.0920 ± 0.001 |
| *R_g_*, (nm) | 4.51 ± 0.4 | 4.11 ± 1.2 | 1.58 ± 0.5 | 4.71 ± 0.4 | 4.64 ± 0.9 |
| *qR_g_* range | 0.65 – 1.10 | 0.71 – 1.11 | 0.47 – 1.27 | 0.75 – 1.27 | 0.67 – 1.26 |
| ***p(r*) analysis** | | | | | |
| *I*(0) (cm^-1^) | 0.4718 ± 0.002 | 0.0657 ± 0.001 | 0.0079 ± 0.0005 | 0.2979 ± 0.002 | 0.0923 ± 0.001 |
| *R_g_* (nm) | 4.88 ± 0.3 | 4.66 ± 0.7 | 3.04 ± 2.5 | 5.18 ± 0.7 | 4.77 ± 0.9 |
| *D_max_* (nm*)* | 16.5 | 15.4 | 11.1 | 19.0 | 17.5 |
| Total estimate from GNOM | 0.70 | 0.88 | 0.50 | 0.68 | 0.47 |
| Quality of fit (χ^2^ / P-value) | 0.92 / 0.55 | 0.93 / 0.54 | 0.90 / 0.78 | 0.90 / 0.81 | 0.97 / 0.81 |
| **Molecular Weight analysis** | | | | | |
| Theoretical MW | 100.45 kDa | | | | |
| MW *I*(0) (kDa) | 36.0 | 34.3 | 654 | 70.0 | 55.9 |
